# Bayesian Trees for Automated Cytometry Data Analysis

**DOI:** 10.1101/414904

**Authors:** Disi Ji, Eric Nalisnick, Yu Qian, Richard H. Scheuermann, Padhraic Smyth

## Abstract

Cytometry is an important single cell analysis technology in furthering our understanding of cellular biological processes and in supporting clinical diagnoses across a variety hematological and immunological conditions. Current data analysis workflows for cytometry data rely on a manual process called *gating* to classify cells into canonical types. This dependence on human annotation significantly limits the rate, reproducibility, and scope of cytometry’s use in both biological research and clinical practice. We develop a novel Bayesian approach for automated gating that classifies cells into different types by combining cell-level marker measurements with an informative prior. The Bayesian approach allows for the incorporation of biologically-meaningful prior information that captures the domain expertise of human experts. The inference algorithm results in a hierarchically-structured classification of individual cells in a manner that mimics the tree-structured recursive process of manual gating, making the results readily interpretable. The approach can be extended in a natural fashion to handle data from multiple different samples by the incorporation of random effects in the Bayesian model. The proposed approach is evaluated using mass cytometry data, on the problems of unsupervised cell classification and supervised clinical diagnosis, illustrating the benefits of both incorporating prior knowledge and sharing information across multiple samples.

## 1. Introduction

Recent advances in high-throughput mass cytometry allow for the measurement of a variety of cellular properties at single-cell resolution (Spitzer and Nolan, 2016). For an individual subject, data can consist of 50 or more marker (or feature) measurements for millions of cells from the subject. This type of data is invaluable for improving our understanding of biological properties such as cellular diversity, as well as playing a key role in the clinical diagnosis of blood cancers such as leukemia and immunodeficiencies such as HIV infection (Wu et al., 2013; Abraham and Aubert, 2016).

A key step in the analysis of such data, in both biological and clinical contexts, is the classification of individual cells into canonical types (e.g., subset populations in lymphocytes such as T cells and B cells). At present the most widely used approach for determining cell types is *manual gating* (Verschoor et al., 2015), based on visual inspection of low-dimensional representations of the data and drawing of bounding boxes or polygons around clusters of cells. The cells that fall within a box are then visualized in another scatter plot involving new dimensions, recursively generating a tree-structure. The selection of particular markers at each node, as well as the placing of the gates, is based on the subjective judgment of the human gater, informed by visual examination of the data being analyzed.

Figure 1 illustrates the manual gating process for cells from one subject. The x and y axes in each subplot correspond to two particular markers selected by the human gater. The leftmost plot is the root node of the tree where the human gater has drawn two polygons (two gates) that determine two subgroups of points. The cells corresponding to each subgroup are then recursively analyzed at the next level of the tree, where additional gates are drawn, and so on. A branch is terminated when the human gater declares a node to be a leaf node, at which point all cells within a particular gate are assigned to a particular cell-type.

**Figure 1:**
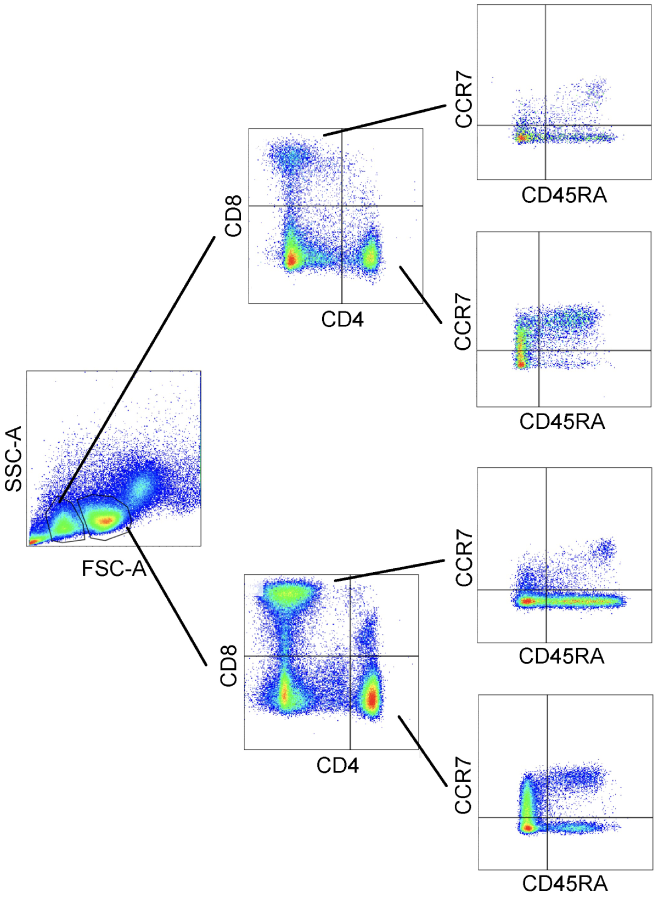
An example of a sequential manual gating procedure, represented via two-dimensional gates at each node in the tree. Each point in a particular plot is an individual cell. Overlaid on each scatter plot are the boundaries of gated regions as determined by a human gater. The x and y axes in each subplot are labeled with the corresponding markers (or features).

Although manual gating is the current default method in practice for analyzing cytometry data, it has a number of significant limitations in terms of its effectiveness and efficiency (Aghaeepour et al., 2013; Verschoor et al., 2015; Saeys et al., 2016). In particular, the method is subjective and heuristic in nature, and hence, is not reproducible in a reliable fashion across samples and studies. In addition, as mass cytometry measurement technology scales to 50 or more features, the limitations of human visual analysis become more apparent and the manual approach is significantly less likely to take full advantage of the high-dimensional measurement space (Chester and Maecker, 2015). Another challenge is presented by the fact that modern cytometry data sets often consist of multiple samples with significant biological and technical variability, e.g., multiple samples per subject over time, samples from multiple subjects in a particular study or group, or samples from different labs or studies. Manual gating is particularly difficult in such contexts, where gating needs to be performed in a consistent and robust manner across multiple samples rather than on a single data sample.

Thus, replacing or augmenting the process of manual gating via the use of automated or semiautomated algorithms is appealing. An algorithm can in principle provide a more reproducible, efficient, and thorough analysis of the data (Qiu et al., 2011). Existing automated approaches, however, have drawbacks that have limited their practical adoption to date in both research laboratories and clinical practice (Kvistborg et al., 2015). In particular, current approaches are unable to incorporate the type of prior knowledge that human gaters use to decide where to draw the bounding boxes and which pair of dimensions to plot next. For instance, a human gater would know that B-cells exhibit high values for marker CD19 and low values for marker CD3 and use this information to guide the cell classification process.

One exception in this context is the recently proposed ACDC algorithm (Lee et al., 2017) that combines prior biological information with mass cytometry data for cell-type classification. The ACDC approach represents prior knowledge in the form of a table relating cell types to markers. The Bayesian approach we describe in this paper is motivated and inspired by this prior work, and in particular by the use of an expert-generated table of prior knowledge to guide the automated gating algorithm.

The algorithm we develop requires (1) a prior knowledge table relating markers and cell-types, and (2) unlabeled multi-dimensional single-cell cytometry data. The output consists of an interpretable hierarchical classification of cells that is consistent with prior biological knowledge. We extend the approach to develop a framework to handle multiple related samples, enabling learning of multiple trees that share common prior knowledge but also reflect differences (via individual random effects) across samples. We evaluate the proposed methodology on real-world mass cytometry data sets. The experimental results demonstrate that the approach produces accurate and interpretable results for both (a) classification of cells into cell types and (b) clinical diagnosis of subjects.

### Clinical Relevance

Through the measurement of a large number of intracellular and cell surface markers at the single cell level, mass cytometry provides the ability to phenotypically and functionally profile individual cells in different disease states. It therefore provides a more effective approach to identify and associate a large number of molecular signatures with heterogeneous clinical outcomes for supporting precision diagnosis of immune system disorders and blood cancers, as well as novel drug development targets. The automated analysis approach proposed in this paper, based on data-driven Bayesian inference, provides a reliable alternative to the current ad hoc approaches to clinical cytometry data analysis, and has the potential to significantly reduce the reliance on subjective manual gating analysis.

### Technical Significance

While our approach shares the same starting point as the afore-mentioned ACDC algorithm, our methodology is different in two important aspects. First, the underlying model in our approach is tree-structured, providing a gating hierarchy that mimics the gating procedure of human gaters, leading to results that are more readily interpretable compared earlier work in this context such as the clustering and random walk approach of ACDC. Second, we cast the problem of learning gating trees in a Bayesian framework, allowing for straightforward development of extensions to the basic model such as the use of random effects to handle multiple data samples in a principled way.

## 2. Methods

Bayesian inference is based on the idea of defining a posterior distribution over quantities of interest (e.g., tree structures) given observed data (e.g., cell-level marker measurements). The posterior distribution is defined via Bayes’ rule, within a proportionality constant, as the product of a prior distribution and a data likelihood. Thus, our Bayesian approach for automated gating consists of three primary components: (1) specification of a prior distribution over gating tree structures, (2) specification of the likelihood of observed data given a particular tree structure, and (3) sampling-based search to find trees that have high posterior probability given the prior and observed data.

### 2.1. Specifying Prior Distributions over Tree Structures

We begin with expert-provided prior knowledge of the same form as used by Lee et al. (2017), namely a table *T*_*C*×*D*_, with *D* markers (features) corresponding to columns and *C* cell-types as rows, as illustrated in the lower part of Figure 2. The table entry for each pair of cell-type and marker is generated based on prior expert knowledge of the expected relationship between the marker and the cell-type. Each entry is specified as *high, low*, or *neutral* (where neutral means unknown or irrelevant), represented as {+1, −1,0}. We assume in our approach (as in Lee et al. (2017)) that the prior information is specified in this tri-valued form, but more fine-grained information (if available) could also be incorporated into the prior. Table 1 provides a specific example of this type of information, a subsample of a table that is used later in the paper in the experimental results section.

**Table 1:** Tabular representation of prior information, i.e., a prior table *T*_*C*×*D*_.

| Cell Types | Markers (dimensions) |  |  |
| --- | --- | --- | --- |
|  | CD4 | CD8 | CD3 |
| Basophils | 0 | -1 | -1 |
| CD4 T cells | +1 | -1 | +1 |
| CD8 T cells | -1 | +1 | +1 |

**Figure 2:**
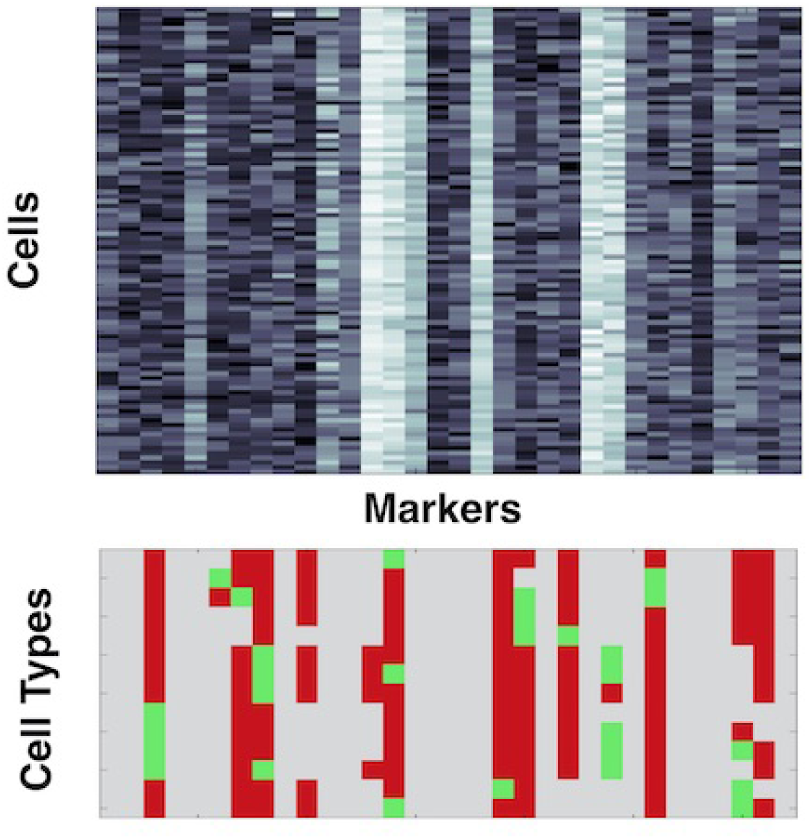
Examples of (a) a subset of mass cytometry data for a subject represented by an *N* × *D* data matrix *χ* (top) with *D* marker measurements (columns) for each of *N* individual cells (rows) from a human subject, with gray-scale indicating marker intensity; and (b) a corresponding prior information table *T*(bottom) of size *C*× *D*, where the *C* rows correspond to known cell types and the *D* columns are markers (same as in the data *χ*). The colors green, red, and gray in the prior table correspond to the cases when cells of a certain type are expected to have a high, low or neutral response to a marker, respectively.

The prior table *T*_*C*×*D*_ can be interpreted as priors on which dimensions a human gater would use to draw a boundary or cut, for a particular marker and for a particular cell-type. In particular, +1 indicates that a cell-type tends have a relatively high response for a particular marker, −1 indicates that a cell-type tends to have a relatively low response for the marker, and 0 indicates that there is no prior knowledge about the relationship between the cell-type and the marker, or that the relationship is not informative with respect to identification of the cell-type. For each marker (corresponding to a column in the prior table) we specify the prior as one of three different Beta densities, depending on the set of +1, 0, −1 values across cell-types for this marker^1^. If the set contains both +1 and −1, the prior on cuts is defined as Beta(*ϕ*_0_, *ϕ*_0_). Setting both Beta parameters to the same value produces a symmetric Beta density with a mode at 0.5, reflecting the fact that there is no strong prior bias for the cut to be low or high. If just +1 (or just 1) is present in the set for the marker, then Beta(*ϕ*_0_, *ϕ*_1_) (or Beta(*ϕ*_1_, *ϕ*_0_)) is used as the prior for cuts for the marker.

In order to mimic the gating process, we also need to specify a prior over the order in which dimensions are selected in a hierarchical fashion. The particular prior distribution that we use is known as a *Mondrian process* (MP). An MP is a flexible recursively-defined Bayesian nonparametric stochastic process in which a finite region is segmented into rectangular partitions (Roy and Teh, 2009). The partitions produced by sampling from an MP result in structures that look like paintings in the style of artist Piet Mondrian. Figure 3 shows a random sample of a partition of a two-dimensional space, from a Mondrian Process prior with a “lifetime” parameter λ_0_ = 1. The Appendix provides additional details on the definition of MP priors.

**Figure 3:**
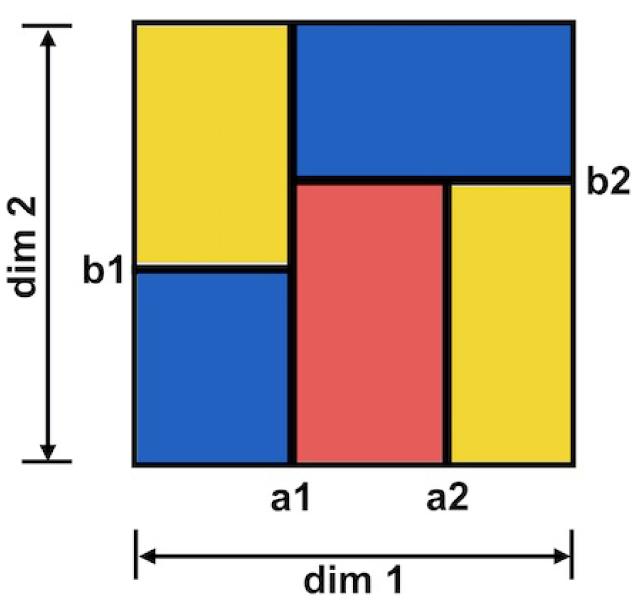
A random sample of a partition of a two-dimensional space from a Mondrian Process prior. The generating process of the above sample is as follows: at the root of the Mondrian tree, a cut was drawn at *a*_1_ in the first dimension, partitioning the entire space into two subspaces; then recursively the two subspaces were further partitioned by random draws from the process corresponding to (dim 2, *b*_1_) and {(dim 2, *b*_2_), (dim 1, *a*_2_)}

A gating tree structure can be sampled from an MP prior by recursively sampling nodes in the tree structure, where each node corresponds to a single marker^2^, drawing a cut for that marker using the Beta priors specified above, and then recursing on the data either side of the cut. The process for sampling a set of nodes (to define a tree structure) is as follows. For any node in the tree (starting with the root node and recursively proceeding from there) a marker 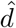 is sampled according to 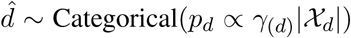^3^, where |χ_*d*_| is the linear dimension (range) of χ_*d*_ and *γ*_(*d*)_ is a scalar taking values from *γ*_0_, *γ*_1_, *γ*_2_ as shown in Table 2, depending on the set of +1/-1/0 values in the corresponding column of the prior information table.

**Table 2:**
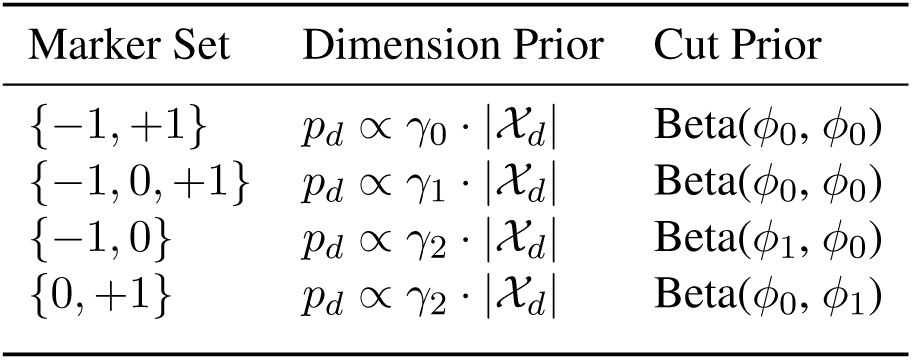
Summary of how the parameters of the priors are defined based on information from the prior table.

Dimensions with both high (+1) and low (−1) values are upweighted by a factor of *γ*_0_, making them more likely to be closer to the root of the tree. This strategy is inspired by the use of information gain to build decision trees in classification algorithms, placing the more discriminative features closer to the root of the tree. Dimensions with high and low values, but also with neutral 0 values, are also upweighted, but to a lesser degree, by a weight *γ*_1_, with *γ*_1_ *« γ*_0_. Dimensions with only one informative label are weighted by *γ*_2_, which is set such that *γ*_2_ *« γ*_1_. After drawing from the Beta, we rescale the cut point appropriately for the dimension, i.e.,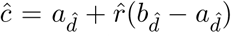 where 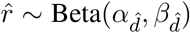. The 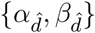 pairs take values from the set {(*ϕ*_0_, *ϕ*_0_), (*ϕ*_1_, *ϕ*_0_), (*ϕ*_0_, *ϕ*_1_)}as shown in Table 2 depending on the marker set of the *d*th column in the prior table.

Lastly, we feed the appropriate sub-tables to two child MPs. Using SQL notation, we perform 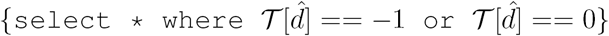 and feed the resulting subtable into the left child (*M*_*<c*_). For the right child, we perform the same query with +1. Thus, upon each recursion,the table contains only the cell types that agree with the cut history. In standard definitions of MPs the sampling process halts when the cost of the sampling process exceeds the “lifetime” parameter λ_0_. However, here we are using informative MPs, so we terminate the sampling process when a table contains exactly one cell type. Figure 4 provides a pseudcode definition of the full procedure for sampling a tree from the MP prior, given a prior information table *𝒯*.

**Algorithm 1.**
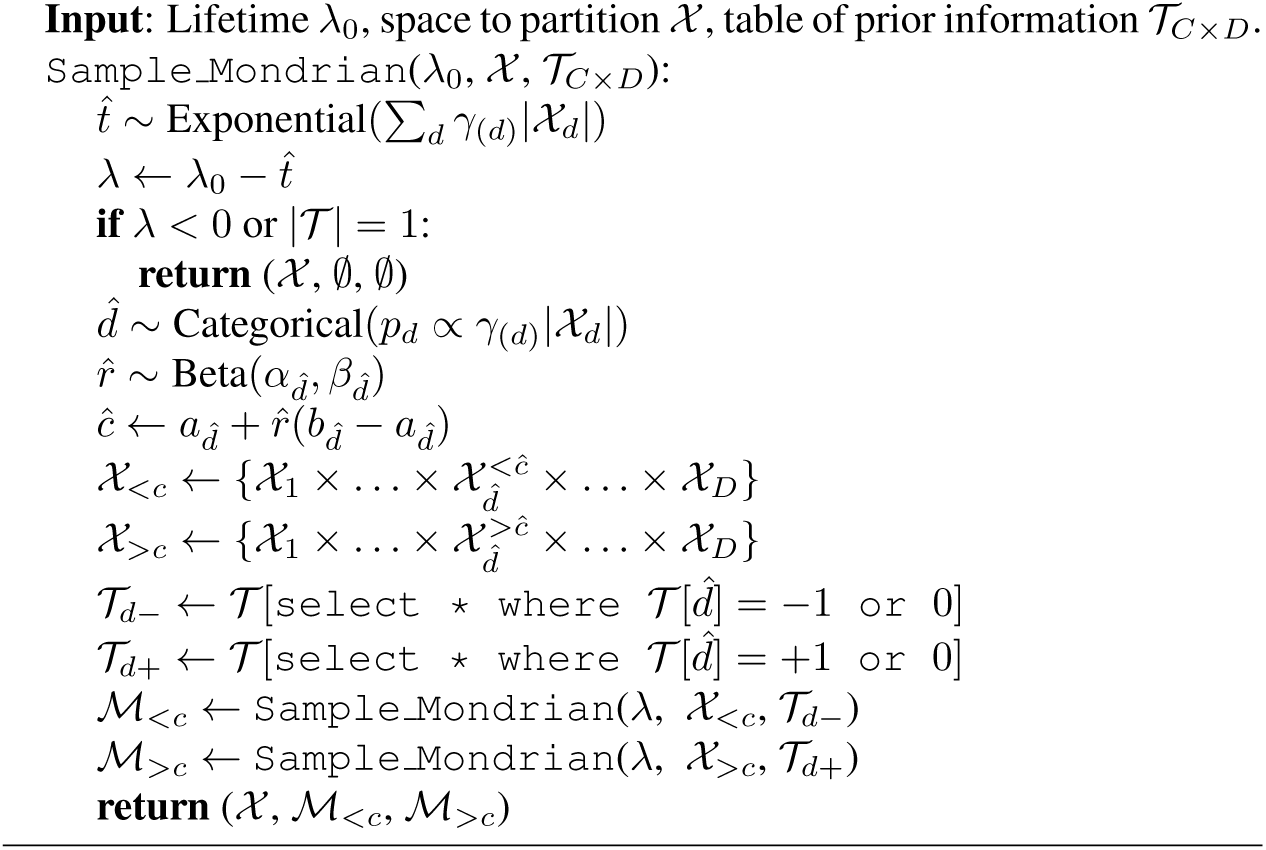
Sampling from a Mondrian Process with Priors

Note that our definition of an MP prior above bears some similarity to method proposed by Wang et al. (2015)’s that incorporates metadata into an MP by adjusting the size of the partitions. However, their approach does not use priors on the cut distributions as we do here.

### 2.2. Modeling Multiple Samples via Random Effects

In using mass cytometry data to learn diagnostic models from labeled data we need to share statistical strength across a collection of samples from different subjects in order to build a model that applies to the entire group. This approach is especially important in clinical applications since we want to exploit all available data to uncover the underlying biological mechanisms. One approach that is commonly used in practice is to pool data across samples (e.g., across subjects), pooling the samples in the healthy group and pooling the samples in group with a disease diagnosis. This approach, however, can result in loss of detail and interpretability at the individual sample level (e.g., for individual subjects).

To accomplish this we can extend the MP prior for a single sample (as described earlier) by placing additive individual-level random effects (REs) on the cut locations. Figure 5 illustrates the basic concept: a global Mondrian template is learned across all data, and individuals or subjects are modeled by using subject-specific random offsets to the globally-defined cuts. Thus, the template is assumed to represent common biological structure while the random effects account for noise due to biological and technical variation across samples. The idea of using random effects for multiple samples in cytometry data analysis has also been pursued by Pyne et al. (2014) in the context of mixture models rather than gating trees.

**Figure 5:**
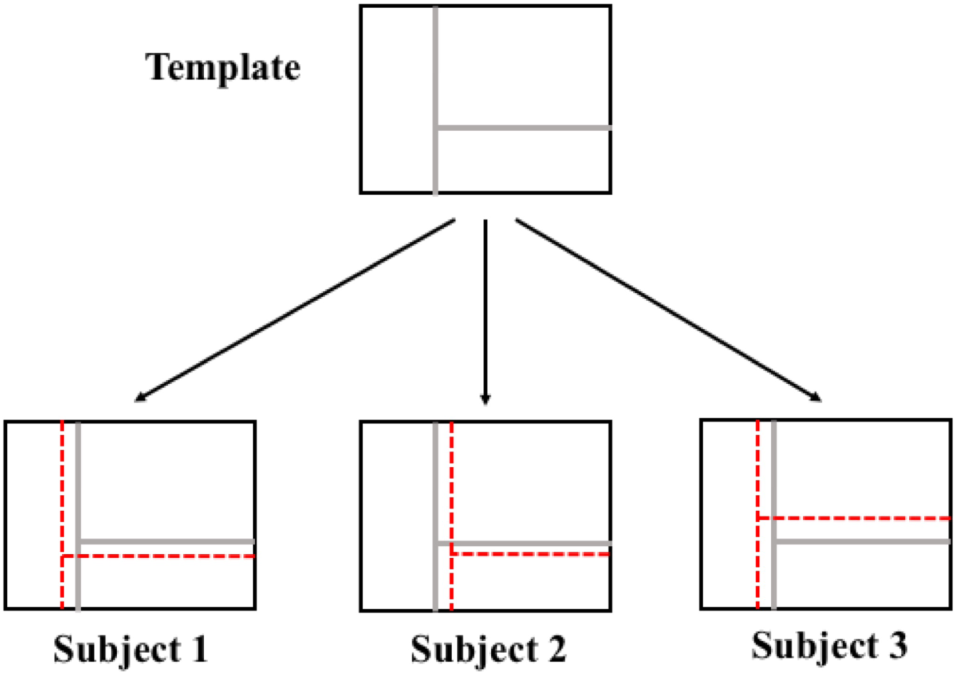
Conceptual illustration of the random effects model in two dimensions, with a template tree on top and individual-level trees of multiple subjects (bottom).

We now define the model formally. Let **X**_*i*_ = {**x**_*i*,1_, *…*, **x**_*i,N*_} (**x**_*i,n*_ ∈ ℝ^*D*^) be *N* data pointsassociated with the *i*th subject (*i* = 1,.., *I*) and, let *χ* be the known or empirical range of all data. We assume there is a global Mondrian tree *ℳ*= {**d**, **c}** with associated dimension indicators **d** and cut locations **c**. A tree for each of the *I* subjects is then created by local Gaussian offsets with covariance matrix Σ. The random effect offset for each subject *i* is denoted as ***ξ*** _*i*_,. Writing the model hierarchically, we have:

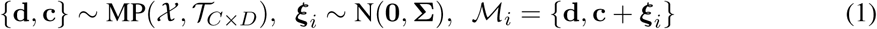

We are interested in learning both the global template and the random effects {***ξ*** _1_, *…*, ***ξ*** _*I*_}The local trees have the same cut orderings as the global model and just the cut locations are different across subjects.

### 2.3. Likelihood of Data given the Model

Having specified the prior over trees, the final step in the modeling process is to define the likelihood of observed data **x**_*i,n*_ given a tree structure for a sample *i*. Given the partitions (ℳ_*i*,1_,….., ℳ _*i, K*_) for the tree ℳ _*i*_, for sample *i*, the likelihood^4^ of a particular data point under a Gaussian likelihood is defined as

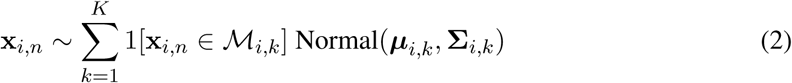

where 1[·] is an indicator function for partition membership, and ***μ***_*i,k*_ ∈ R^*D*^, **Σ**_*i,k*_ ∈ R^*D*×*D*^ are the parameters of the Normal distribution associated with partition *M*_*i,k*_. Intuitively, the model can be thought of as a Gaussian mixture in which the mixture weights are determined by the local tree. If it were possible to marginalize out the tree, the likelihood would be 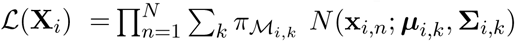 where *π* _*Mi,k*_is the probability that **x**_*i,n*_ is contained in the *k*th partition. As the number of partitions is random, we have a model similar in spirit to the Dirichlet process mixture model (Neal, 2000), with the difference that the membership probabilities are determined by the MP’s tree structure instead of the Dirichlet process’ preferential attachment procedure (Ferguson, 1973).

### 2.4 Learning Trees and Cuts given Data

Previous work (Roy and Teh, 2009; Wang et al., 2015) performed posterior inference on trees and cutpoints via Markov chain Monte Carlo (MCMC) sampling, with sampling proposals consisting of drastic changes to the segmentation structure (translations, rotations, re-drawing, etc. of the partitions). While this approach is suitable for two-dimensional problems, it is computationally impractical for the high dimensional trees we need for cytometry analysis. Instead, we perform approximate inference by sampling tree structures from the prior rather than sampling trees by conditioning on the data. While this does not correspond to exact posterior inference, we found it worked well in practice, likely due to the fact that the prior is based on strong biological knowledge. Given the sampled trees, we then run MCMC on just the cut locations using local Gaussian proposals.

In terms of computational costs, the time complexity of sampling a single tree structure *T* from the prior distribution is *O*(*KD*), where *K* and *D* are the number of cell types and markers respectively. Given a tree structure *T* the complexity of proposing and sampling one set of cutpoints *C* is *O* (*ND*^2^ + *KD*^3^), where *N* is the number of cells in the sample. Thus for each MCMC iteration,the amortized complexity is *O*(*ND*^2^ + *KD*^3^).

In the results for cell classification described in the remainder of the paper, we draw 50 sampled trees *T’* from the MP prior and set the informative priors such that *γ*_0_ = 1000, *γ*_1_ = 100, *γ*_2_ = 1, and *ϕ*_0_ = 5, *ϕ*_1_ = 2. We then run 50 MCMC chains (one per tree *T ’* from), where each MCMC chain consists of 3000 iterations. Each iteration is a draw *C’* from of cutpoints conditioned on the corresponding tree *T ’ C’* is generated from a Gaussian distribution centered at the current sample *C* with covariance matrix 0.1 times the identity matrix. The last 20 samples from each chain (20× 50 = 1000 samples overall) are then used for prediction. In terms of the number of MCMC iterations required to obtain good results, the encoding of relevant scientific knowledge into the MP prior results in an informative prior distribution that is much closer to the true posterior compared to typical non-informative priors used in MPs. This informative prior makes inference much less computationally intensive than with a non-informative prior, allowing the MP to be used in much higher dimensions than in previous work. For the disease diagnosis results (with multiple samples) reported later in the paper we used the same procedure as above except that we draw 32 chains and 1000 samples for each chain. Random effect offsets are sampled from a zero-mean Gaussian distribution with a covariance matrix defined as 0.05 times the identity matrix.

For illustration, Figure 6 shows an example of the upper portion of an MP tree learned using the procedure above for single-cell mass cytometry data from a blood sample for a single individual (details about this particular data set are described in the next section). The tree reflects prior knowledge (in terms of tree structure) in addition to being faithful to the data (in terms of cut locations). It is also directly interpretable in that it mimics a typical human gating procedure with one-dimensional cuts, and leads to cell type classifications (at the leaf nodes) that can be easily understood in terms of thresholds on the measurement markers.

**Figure 6:**
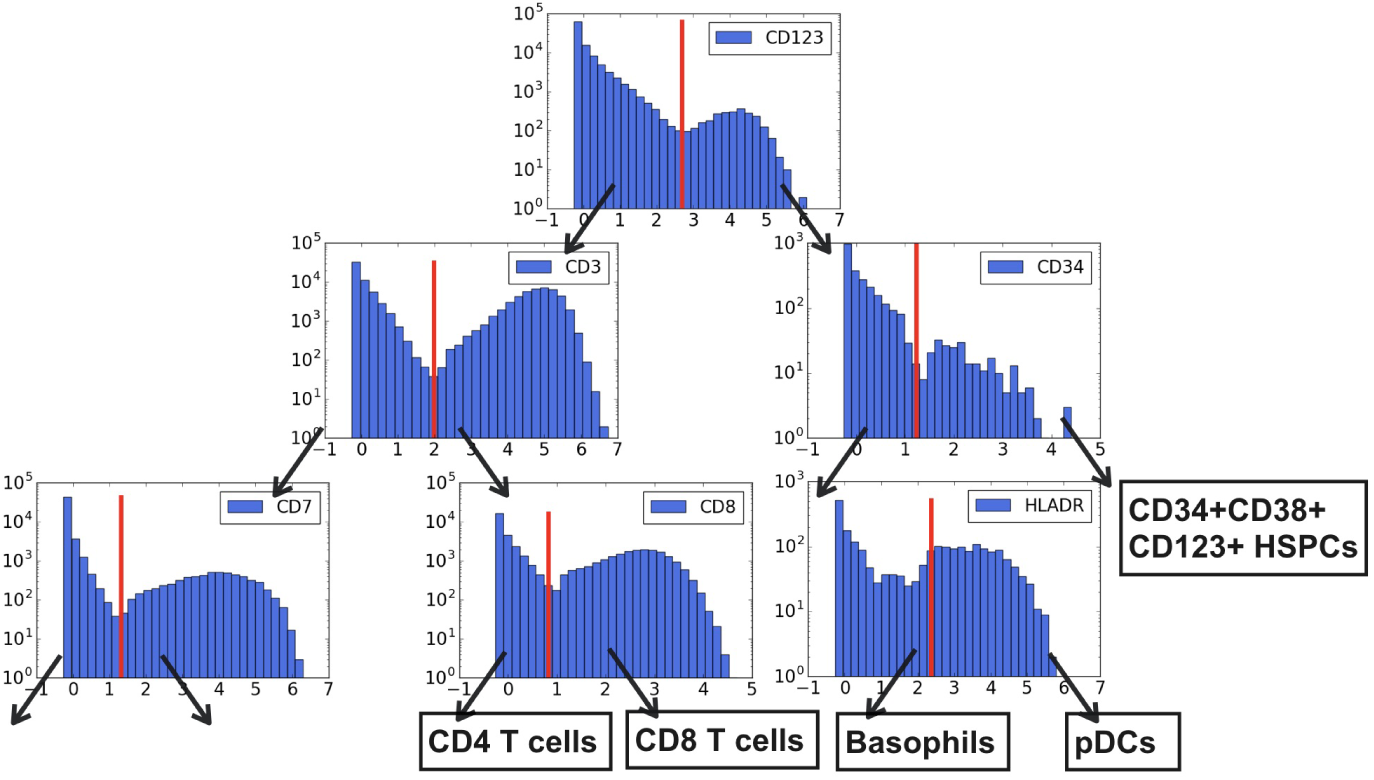
The upper portion of a tree structure for the MCMC sample with the highest joint probability over data, cut points, and tree structures for the AML data set. Red lines denote sampled cuts, and arrows denote the path taken by cells that fall on the left or right side of the cut. The black rectangles denote cell type classifications.

Figure 7 shows examples of posterior samples (in particular dimensions) for the data sets (AML and BMMC) used in the experimental results section. Each blue line represents a sampled cut and each black dot represents a cell for the two plotted markers (CD4 vs CD3 for AML, CD4 vs CD8 for BMMC). The figures show 100 posterior MCMC samples for two markers on each data set. For AML (top), 97 out of 100 posterior MCMC samples drew the first cut on CD3, because in the prior information table for the AML data the marker label sets of CD3 and CD4 are {-1, +1} and{-1, 0, +1}. Based on the prior distributions the probability of CD3 is upweighted by the factor 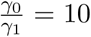 compared to CD4. For BMMC (bottom) 62 out of 100 MCMC samples place the first cut on CD4, while the other 38 MCMC samples place the first cut on CD8. This is because both of the marker label sets of CD4 and CD8 are {1, +1}, and thus, the probability of drawing a cut from each dimension is proportional to its scale. These types of plots are similar to the displays currently used in manual gating of cytometry data and, hence, are directly interpretable to cytometry experts. This type of information could for example be used to assist human gaters via an interactive visual interface for semi-automated gating.

**Figure 7:**
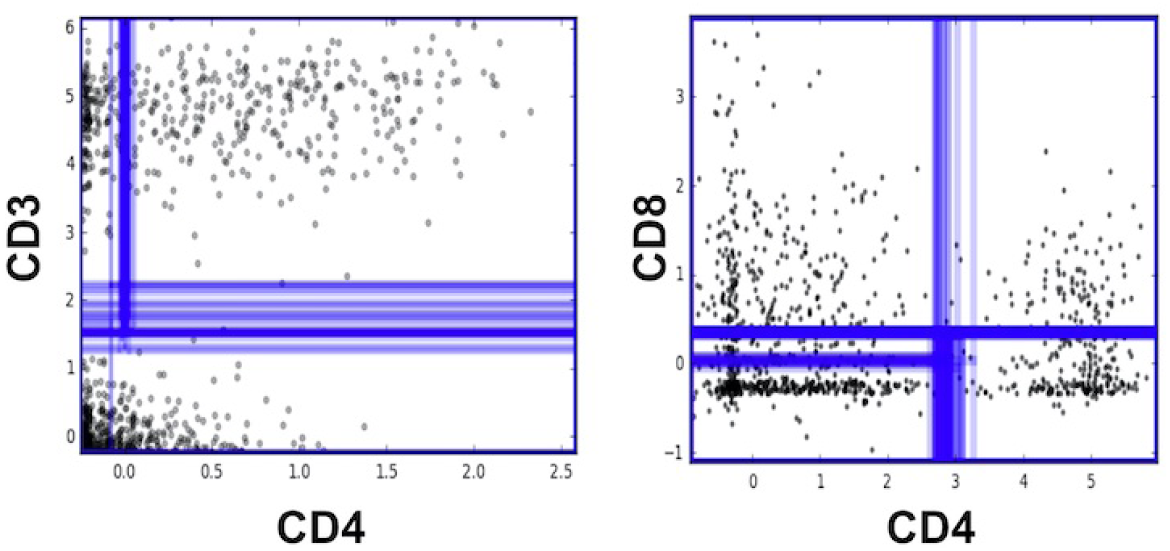
Examples of posterior samples of cutpoints for pairs of markers for the AML (left) and BMMC (right) data sets.

We also investigated the sensitivity of our sampling procedure to changes in the parameters for the Beta distribution, *ϕ*_0_, *ϕ*_1_, and found that the inference results are not sensitive to changes as long as the prior distribution is reasonably flat. We found, however, for the Gamma prior parameters, *γ*_0_, *γ*_1_ and *γ*_2_, that it is important to ensure that *γ*_2_ «*γ*_1_ «*γ*_0_ so that markers which are more informative in terms of classification are prioritized to appear higher in the tree. Otherwise the process of sampling tree structures will generate many implausible trees and the MCMC search will be computationally inefficient.

## 3. Results: Classifying Cell Types

Although our model requires no cell-level labels, we can perform cell type classification by using the prior knowledge table combined with the MP posterior distribution. As illustrated in Figure 6, given an MP tree, each cell takes a path from the root node to one of the leaf nodes by taking the left or right side of the cut at each step. We find the MP partitions that obey the table constraints for each type. For example, CD8 T cells are assigned to the partition on the high side of the cut on CD8, high side of CD3, etc. If two or more cell types are assigned to the same partition we classify cells in this partition randomly to these cell-types. For the 1000 MCMC samples that we obtain, each of them can be viewed as a classifier of cells into types. For cell classification we use an ensemble approach and take the majority vote of the 1000 predictions.

We evaluated our approach on mass cytometry measurements of human cells and prior information tables used in Lee et al. (2017). The samples are from two public benchmark data sets: acute myeloid leukemia (AML) (Levine et al., 2015) and bone marrow mononuclear cells (BMMC) (Bendall et al., 2011). The AML data set consists of 32 markers and has been manually gated into 14 cell types. The BMMC sample has 13 markers and 19 cell types (cell types without any prior information were removed).

Table 3 reports cell type classification accuracy of our method (MP) against two baselines that represent two extremes: a Gaussian mixture model (GMM) approach that has no prior knowledge, and an approach based only on sampling from the prior (MP-Prior) that ignores the data. For each of the methods the partitions or clusters derived by the method are matched up (automatically for the MP approaches, and manually for GMM) to known human labelings of cells into cell types that are known for these data sets from manual gating. We see that our model significantly outperforms both.

**Table 3:** Cell classification accuracy on AML and BMMC data sets for different methods.

|  | AML | BMMC |
| --- | --- | --- |
| **Methods without Cell-Level Labels** |  |  |
| MP (Proposed Method) | 96.9% | 92.3% |
| MP-Prior | 61.5% | 85.6% |
| ACDC | 98.2% | 93.7% |
| **Methods requiring Cell-Level Labels** |  |  |
| GMM | 86.1% | 84.1% |
| Phenograph | 95.1% | 95.0% |

For comparison with other cell-level cytometry data analysis methods, we also included the results from two other classification algorithms, ACDC (Lee et al., 2017) and Phenograph (Levine et al., 2015). ACDC uses the same prior information table as we use in our MP approach, to identify and classify a subset of cells as landmark cells, and the remaining cells are then classified into canonical cell types via a random walk approach. For the phenograph clustering method, the data are first clustered using community detection and then each cluster is assigned to a manually-gated cell-type. The accuracy of our method is comparable with both ACDC and Phenograph on both of the data sets. A key point, however, is that the MP approach is more directly interpretable to cytometry experts than the ACDC or Phenograph methods since it mimics the hierarchical manual gating process that is widely used in clinical applications involving cytometry data. In addition, the MP method is able to achieve relatively accurate performance using general prior knowledge without requiring any manual labeling of individual cells per data set—this becomes particularly relevant with dealing with multiple samples (discussed below).

Additional insights can be gained by comparing the algorithm’s predictions with the true labels. For the AML data set, Figures 8 and 9 show t-SNE plots and confusion matrix (respectively) for predicted and true labels. Both the confusion matrix and the t-SNE plots demonstrate that the algorithm is producing high-fidelity labels that closely match the detailed cell-level labels produced by human gating.

**Figure 8:**
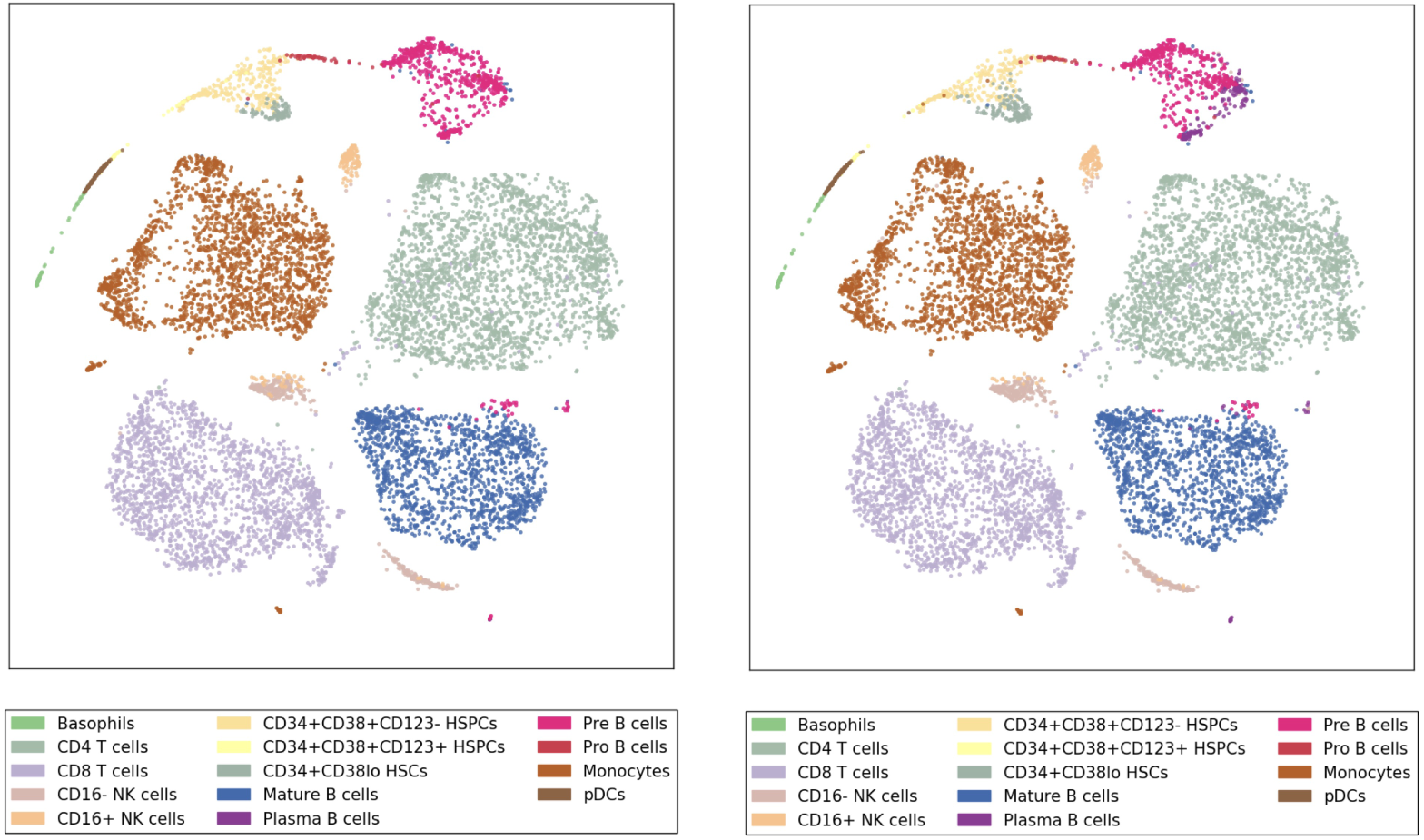
t-SNE map for AML data with true labels (left) and with predicted labels (right).

**Figure 9:**
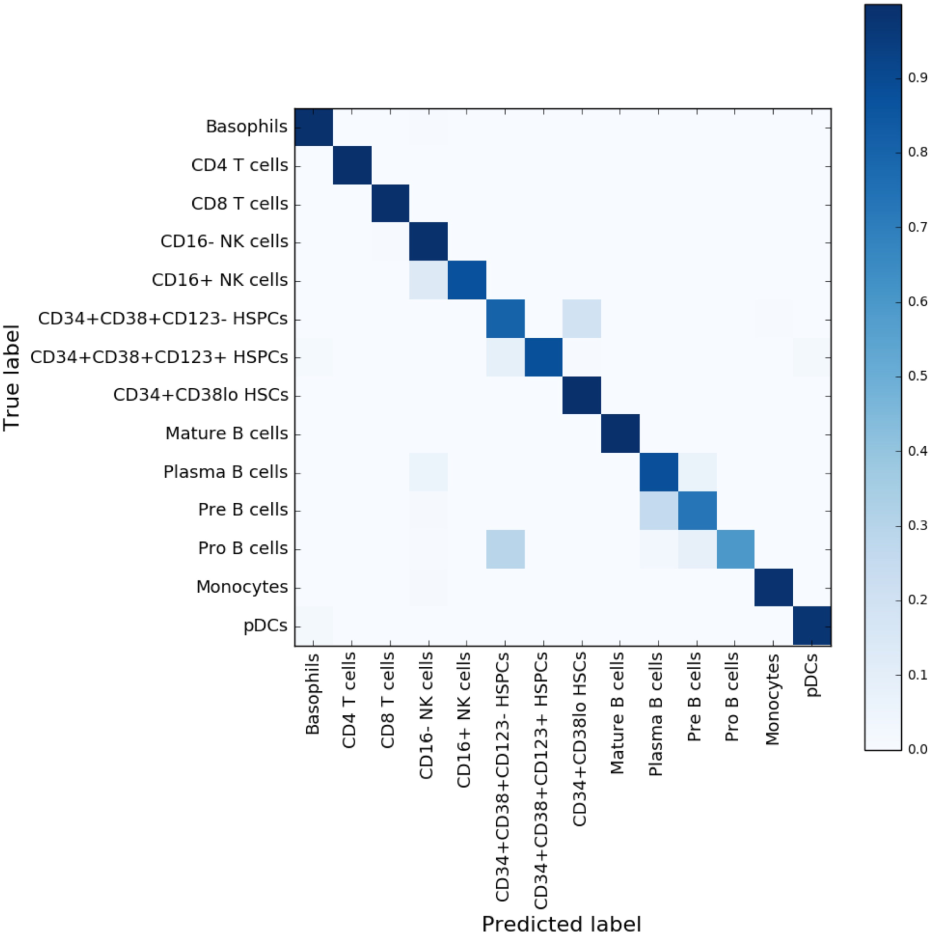
Confusion matrix for AML data: comparing true labels (rows) with predicted labels (columns). Each entry in the table is color-coded based on the fraction of cells that truly belong to a cell-type (row) that were predicted to belong to a cell-type (column).

## 4. Results: Disease Diagnosis

In clinical diagnosis the goal is to classify (diagnose) whether a subject has a particular disease or not given one or more samples for that subject. The problem is challenging because of biological variability across individuals, different numbers of cell measurements per individual, and the lack of large labeled data sets (in terms of number of individuals). Our MP-RE approach is well-suited to handle these issues. To handle variability across individuals, while still sharing common information, the MP-RE can share information across individuals within each of the healthy and non-healthy groups while still allowing for individual variability via the random effects.

We evaluated the our approach on a well-known acute myeloid lymphoma (AML) mass cytometry data set from Levine et al. (2015) consisting of cell-level data with 16 markers for 5 healthy subjects and 16 subjects diagnosed with AML. Prior knowledge was obtained from the same expert tables provided for these markers by Lee et al. (2017). The prior knowledge is common across both groups in terms of which markers are relevant to which cell-types. Note that we have supervised information (healthy versus AML labels) at the level of subjects, but there are no labels available at the cell level for these data sets.

To classify individuals into two groups we proceeded as follows. Two different MP-RE models were fit during training, one to the samples in the healthy group and one to the samples in the AML group. The MP-RE constructs a template Mondrian tree for each of the two groups (using the MCMC procedure described earlier) where each template tree models the overall characteristics of subjects in the group. Each subject’s individual Mondrian tree contains subject-specific offsets to the cuts (random effects) relative to the template Mondrian tree.

To classify a new sample of cells, we fit two Mondrian trees with random effects to the sample, one where we estimate an MP-RE tree for the sample using the healthy Mondrian template and the other with the AML template. This results in a partitioning of the cells in the sample into *K* cell-types, from the “perspective” of each of the healthy and AML templates. We compute the proportion of cells assigned to each of the *K* cell-types for each tree, resulting in two *K*-dimensional vectors, which we concatenate to create a final feature vector for prediction per sample.

We used leave-one-out cross-validation to evaluate classification accuracy, leaving out each individual sample, fitting a standard logistic regression model to the cell-type proportions for the other samples, and using the logistic model to predict the diagnosis of the left-out sample. The proportions of cell-types for each sample were estimated for each sample as described earlier. We compared our MP-RE method with two other baseline approaches. The first baseline is where we globally pooled all of the cells from each sample within each group (healthy and AML) to learn two MP trees, where the trees use the prior knowledge but there are no individual random effects (due to pooling). Each MP tree classifies cells of the test sample into cell types by partitioning the data space. The second baseline was another pooled approach (all cells in all samples per group) where k-means was fit to each pooled set of cells. For a test sample, cells are assigned to cell types of the nearest kmeans cluster center. Both baselines were evaluated using the same leave-one-out strategy as the MP-RE method, where the feature vector for the test sample was computed by passing it into the cell-type classifier learned on pooled data of each group.

MP-RE and Global MP predicted the correct class label for all 21 samples, while k-means produced 20 out of 21 correct predictions. While the Global MP produced correct predictions, the MP-RE model can provide a more nuanced and useful interpretation of the data. For example, we further analyzed the distribution of the sizes of the estimated random effects within the two groups in Figure 10 (left). The variability of random effects in the AML group is systematically greater than that of the healthy group. This observation is consistent with medical knowledge: more within-group variability is expected in marker measurements of AML patients than healthy patients, given that the AML patients may be at different stages of the progression of AML. Figure 10 (right) shows an example of the random effects (the gap between red line and green line) for the first cut of MP trees for subjects H1, H2, H5. With an additive random effect fit to data, the cut location of H5 moved towards right, because its upper component contains more data points compared to H1 and H2.

**Figure 10:**
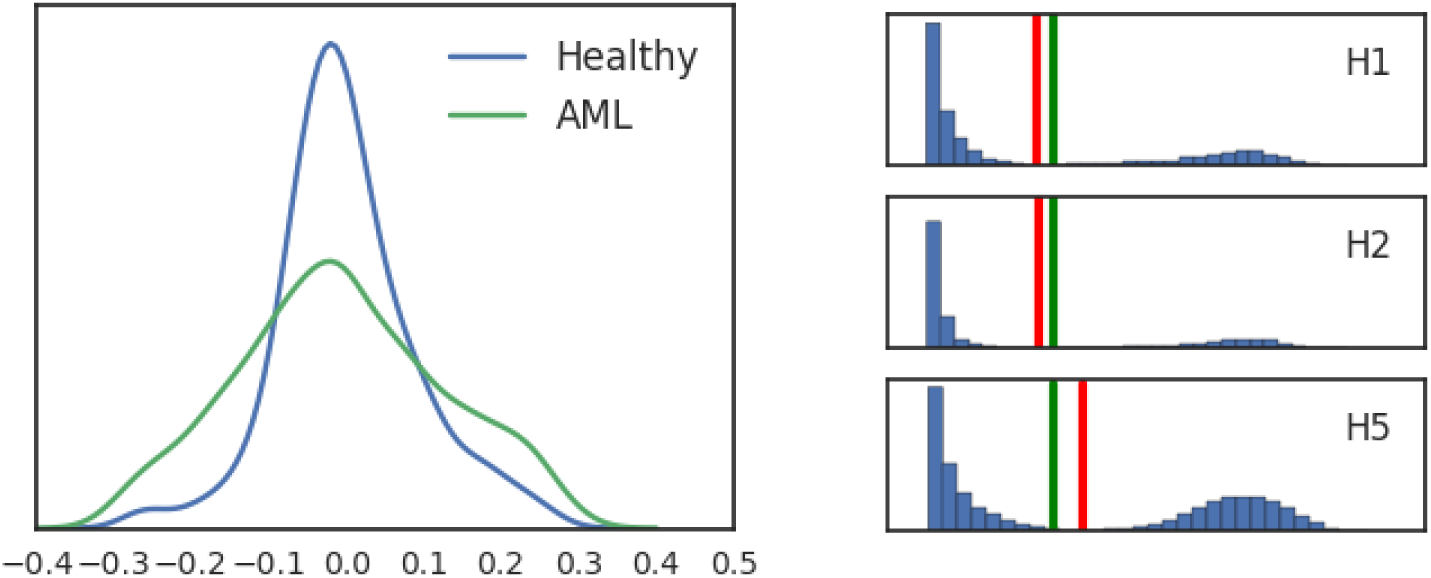
Results from the MP-RE model on healthy and AML subjects. (Left) Density estimate of estimated random effects (relative to the scale of the corresponding dimension), in the healthy group and in the AML group, across all subjects in each group and across all MCMC chains. (Right) Estimated random effects for three healthy subjects H1, H2, H5 for marker dimension CD123. The green line represents the cut in the template MP, which is shared across all subjects. The red line represents the individual cut adjusted from the template based on subject’s data. (Best viewed in color).

## 5. Conclusions

In this paper we proposed a Bayesian framework for automated gating to classify cells into cell-types, combining prior knowledge with observed cytometry data. Our approach to representing prior information is motivated by the recent work of Lee et al. (2017), who proposed the use of prior tables relating pairs of cell-types and markers as part of the ACDC algorithm. However, ACDC uses multiple algorithmic steps (unsupervised clustering, random walks on a nearest neighbors graph) in contrast to the more interpretable tree-structured model we propose here.

Our empirical results demonstrate that the proposed Bayesian approach is able to use prior knowledge in an effective manner to discover interpretable tree structures that characterize biologically meaningful cell types, and to enable automated clinical diagnosis using data sets from multiple individuals. There are a number of potential directions for future work. We focused on axis-aligned gating boundaries that threshold along specific dimensions, but manual gating can also allow for general polygon segmentations. Such an extension would increase the model’s flexibility but performing inference would be computationally complex. Other possible extensions involve the use of partial prior knowledge, altering the emission model to non-Gaussian densities, modeling multiple dependent samples from the same subject over time, and utilizing a fully Bayesian approach in which hyperpriors for random effects are learned (e.g., for different groups) rather than being fixed.

## Appendix: Definition of the Mondrian Process (MP) Prior

For completeness we include a brief formal definition of the MP prior—for a more thorough discussion see (e.g.) Balog et al. (2016). Define an axis-aligned box to be a product space of *D* bounded intervals *χ*_*d*_ = [*a*_*d*_, *b*_*d*_] with length |*χ*_*d*_ |= *b*_*d*_ -*a*_*d*_: *χ*={ *χ* _1_ ×*…* × *χ*_*D*_.} Define a *Mondrian process* (MP) with a lifetimeλ _0_ and on a space *χ* as MP(λ _0_,. *χ)* The process proceeds by first drawing an exponential random variable 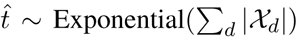. If 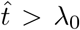 the process halts and returns *χ* without any partitions. If 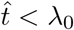 a dimension 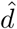 is drawn proportionally to its length (*p*_*d*_ ∞ |*X*_*d*_*|*) and then a cut location 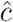 is drawn according to 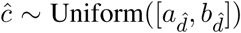. In other words, the space is partitioned by the cut into two new spaces 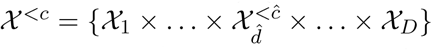 and 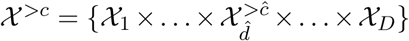 Two child processes 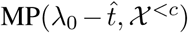 and 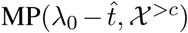 are then spawned and the process recurses with a decreased lifetime 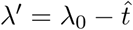 and on the subdomains *χ’*^*<c*^ and *χ”* = *χ*^*>c*^. This recursion arises from the elegant self-consistency property of MPs: further cuts to any partition are themselves drawn from an MP with a lifetime and domain properly inherited from the parent process.

## Footnotes

1 The Beta density *f* (*x*) = (*x*; *α, β*) is a flexible prior density for values *x* ∈ [0, 1] specified by two parameters *α* > 0, *β* > 0.

2 The tree structure defined in this manner corresponds to single-dimensional histogram gates rather than two-dimensional scatter plot gates.

3 Categorical(*·*) is a probability distribution over *d* ∈ {1, …, *D*} where *D* is the number of markers.

4 Densities other than Gaussian could also be used for the data likelihood—we chose the Gaussian model for simplicity.

